# Dose-dependent response to infection with SARS-CoV-2 in the ferret model: evidence of protection to re-challenge

**DOI:** 10.1101/2020.05.29.123810

**Authors:** Kathryn A. Ryan, Kevin R. Bewley, Susan A. Fotheringham, Phillip Brown, Yper Hall, Anthony C. Marriott, Julia A. Tree, Lauren Allen, Marilyn J. Aram, Emily Brunt, Karen R. Buttigieg, Breeze E. Cavell, Daniel P. Carter, Rebecca Cobb, Naomi S. Coombes, Kerry J. Godwin, Karen E. Gooch, Jade Gouriet, Rachel Halkerston, Debbie J. Harris, Holly E. Humphries, Laura Hunter, Catherine M. K. Ho, Chelsea L. Kennard, Stephanie Leung, Didier Ngabo, Karen L. Osman, Jemma Paterson, Elizabeth J. Penn, Steven T. Pullan, Emma Rayner, Gillian S. Slack, Kimberley Steeds, Irene Taylor, Tom Tipton, Stephen Thomas, Nadina I. Wand, Robert J. Watson, Nathan R. Wiblin, Sue Charlton, Bassam Hallis, Julian A. Hiscox, Simon Funnell, Mike J. Dennis, Catherine J. Whittaker, Michael G. Catton, Julian Druce, Francisco J. Salguero, Miles W. Carroll

## Abstract

In December 2019 an outbreak of coronavirus disease (COVID-19) emerged in Wuhan, China. The causative agent was subsequently identified and named severe acute respiratory syndrome coronavirus 2 (SARS-CoV-2) which rapidly spread worldwide causing a pandemic. Currently there are no licensed vaccines or therapeutics available against SARS-CoV-2 but numerous candidate vaccines are in development and repurposed drugs are being tested in the clinic. There is a vital need for authentic COVID-19 animal models to further our understanding of pathogenesis and viral spread in addition to pre-clinical evaluation of candidate interventions.

Here we report a dose titration study of SARS-CoV-2 to determine the most suitable infectious dose to use in the ferret model. We show that a high (5×10^6^ pfu) and medium (5×10^4^ pfu) dose of SARS-CoV-2 induces consistent upper respiratory tract (URT) viral RNA shedding in both groups of six challenged animals, whilst a low dose (5×10^2^ pfu) resulted in only one of six displaying signs of URT viral RNA replication. The URT shedding lasted up to 21 days in the high dose animals with intermittent positive signal from day 14. Sequential culls revealed distinct pathological signs of mild multifocal bronchopneumonia in approximately 5-15% of the lung, observed on day 3 in high and medium dosed animals, with presence of mild broncho-interstitial pneumonia on day 7 onwards. No obvious elevated temperature or signs of coughing or dyspnoea were observed although animals did present with a consistent post-viral fatigue lasting from day 9-14 in the medium and high dose groups. After virus shedding ceased, re-challenged ferrets were shown to be fully protected from acute lung pathology. The endpoints of URT viral RNA replication in addition to distinct lung pathology and post viral fatigue were observed most consistently in the high dose group. This ferret model of SARS-CoV-2 infection presents a mild clinical disease (as displayed by 80% of patients infected with SARS-CoV-2). In addition, intermittent viral shedding on days 14-21 parallel observations reported in a minority of clinical cases.

## Introduction

Coronaviruses are positive sense, single stranded RNA viruses belonging to the family Coronaviridae^1^. These viruses can infect a range of animals, including humans and usually cause a mild respiratory infection, much like the common cold. Two highly pathogenic coronaviruses have emerged in the human population in the last 20 years; severe acute respiratory syndrome (SARS) CoV and middle eastern respiratory syndrome (MERS) CoV. SARS-CoV infected approximately 8,000 people worldwide with a case fatality rate (CFR) of 10%, while MERS-CoV has infected approximately 2,500 people with a CFR of 36% ^2^.

In December 2019 several pneumonia cases of unknown cause emerged in Wuhan, Hubei, China. Deep sequencing analysis from lower respiratory tract samples from patients indicated the cause to be a novel coronavirus^3^. The causative agent of this novel coronavirus disease (COVID-19) was identified as SARS-CoV-2. As of May 22^nd^ 2020, there have been over 4,893,000 confirmed cases reported worldwide, including over 323,000 deaths, in over 180 countries^4^. The global mortality rate is yet to be determined. Approximately 80% of patients display only mild symptoms, with approximately 14% displaying severe symptoms such as dyspnoea and low blood oxygen saturation. Around 6% of cases become critical, with respiratory failure, septic shock and/or multiple organ failure^5^. There is an urgent need to develop suitable animal models to evaluate antivirals or vaccine candidates against SARS-CoV-2.

Ferrets have been used extensively to model the disease caused by influenza virus^6-12^ infection as well as human RSV^13,14^, mumps virus^15^, Ebola virus^16,17^ and Nipah virus^18,19^. Due to the presence of ACE2, the virus receptor, on cells of the ferret respiratory tract, these animals were developed an as effective model for SARS-CoV^20-23^. The ferret has been shown to shed detectable virus from its URT as well as exhibiting comparable clinical symptoms associated with milder cases of the infection^21^ and shown similar pathology in the lung to that observed in humans^22^. SARS-CoV-2 spike protein has been shown to exhibit many similarities in its amino acid sequence and protein structure to the receptor binding domain of SARS-CoV^24^ and also utilises ACE2 for cell entry ^25^, suggesting ferrets would be a potential model for COVID-19.

To understand if ferrets are a suitable model for SARS-CoV-2 infection we challenged animals intranasally with a range of titres of SARS-CoV-2 (5×10^2^, 5×10^4^ and 5×10^6^ pfu) in 1ml volume. The purpose was to characterise the most suitable challenge dose for use in future studies, to aid understanding of the kinetics of viral pathogenesis and the immune response following infection and to facilitate the evaluation of treatments and vaccines against SARS-CoV-2.

## Results

### Study Design

Ferrets were challenged intranasally with 1ml of Victoria/1/2020^26^ SARS-CoV-2 at three different titres representing a high, medium and low dose (**Table 1**). A high titre stock of challenge virus was prepared (passage 3), and quality control sequencing showed it was identical to the original stock received from the Doherty Institute and did not contain a commonly reported 8 amino acid deletion in the furin cleavage site^27^. Following the initial challenge, a re-challenge with the high dose (5×10^6^ PFU) took place at day 28 post challenge (pc). The four (two per group) remaining ferrets in groups 2 and 3 were re-challenged via the same, intranasal, route and 1ml volume alongside a control group of two naïve control ferrets (group 5).

**Table 1.** Experimental Animal Groups.

| Group |  | Number of Animals | Initial Challenge Virus Titre (pfu/ml) | Re-challenge Virus Titre (pfu/ml) | Euthanasia/ challenge days |
| --- | --- | --- | --- | --- | --- |
| 1 | High | 6 | $5 \times 10^6$ | | 1 ferret euthanised at day 3, 5, 7, 14 pc<br>2 ferrets euthanised day at 21 pc |
| 2 | Medium | 6 | $5 \times 10^4$ | $5 \times 10^6$ | 1 ferret euthanised at days 3, 5, 7, 14 pc<br>2 ferrets re-challenged at day 28 pc<br>1 ferret euthanised at days 33 & 36 pc |
| 3 | Low | 6 | $5 \times 10^2$ | $5 \times 10^6$ | 1 ferret euthanised at days 3, 5, 7, 14 pc<br>2 ferrets re-challenged at day 28 pc<br>1 ferret euthanised at days 33 & 36 pc |
| 4 | Naïve Sentinel | 2 | PBS |  | 2 ferrets euthanised at day 20 pc |
| 5 | Naïve Control | 2 | | $5 \times 10^6$ | 1 ferret euthanised at days 33 & 36 pc |

A total of 22 ferrets were distributed across 5 groups. All inoculations were performed intranasally with 1ml of fluid. N/A, not applicable, pc, post-challenge.

### Viral Shedding following challenge

Viral RNA was detected in the nasal wash of 6/6 ferrets in the high dose group from day 1 pc and continued to be detected at varying levels until day 20 pc (**Fig. 1a**). The peak in viral RNA shedding was seen between day 2 and 4 pc for all ferrets in the high dose group. Following a decline in viral RNA (2/2 animals) to below the limit of quantification at day 13 pc, an increase was seen at days 16 and 18, with a measurement in viral RNA for one ferret (4.75×10^4^ copies/ml) just above the limit of quantification at day 16 pc and a viral load of 1.1×10^6^ copies/ml in the other ferret at day 18 pc. Both Group 1 survivors were euthanised on day 21 at which point no viral RNA was detected in their nasal washes.

In the medium dose group 6/6 ferrets also had detectable viral RNA in nasal washes from day 1 pc. The peak of viral RNA shedding was more variable in the medium dose group, with some ferrets peaking at days 2 to 3 pc (4/6) and others peaking at days 5 to 6 pc (2/6). A decline was then seen until day 11 pc where viral RNA levels fell below the limit of quantification, but viral RNA was still detected. By day 16 no more viral RNA was detected. Quantifiable viral RNA was only found in the nasal wash of 1/6 ferrets in the low challenge dose group. This ferret was euthanised on day 5 pc. No other ferrets in the low dose group were found to shed quantifiable viral RNA in their nasal wash.

**Figure 1.**
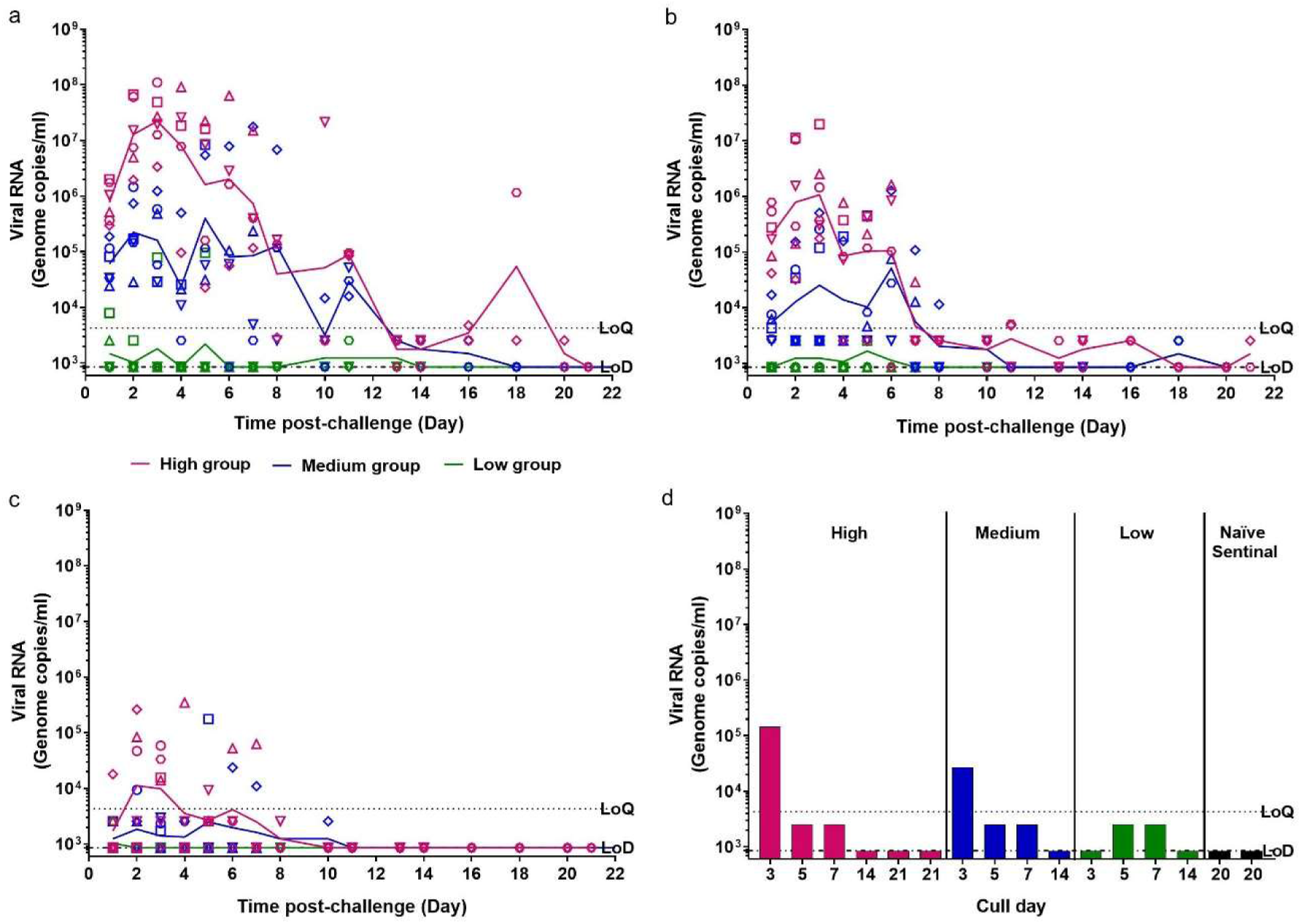
Viral RNA Shedding. Nasal washes and swabs were collected at days 1 to 8, 10, 11, 13, 14, 16, 18 & 20 pc for all virus challenged groups. Viral RNA was quantified by RT-qPCR. No viral RNA was detected in any samples taken from the naïve sentinel ferrets (data not shown). (**a**) Nasal washes (**b**) Throat swabs (**c**) Rectal swabs (**d**) Bronchoalveolar lavage collected at necropsy (numbers indicate day post challenge the ferret was euthanised). Points show values for individual animals, lines show group geometric means. The dashed horizontal lines show the lower limit of quantification (LoQ) and the lower limit of detection (LoD).

A similar trend in the titre of viral RNA detected in nasal wash samples was observed in the throat swabs samples during the first week after challenge (**Fig. 1b**). The amount of viral RNA detected in the throat swab samples of ferrets in the high dose group (6/6) peaked at day 3 pc. In contrast, however, detection of viral RNA in throat swab samples was less prolonged than in the nasal passage, with no quantifiable viral RNA detected past day 11 pc.

Detection of viral RNA in the rectal swabs was found to be variable across the different dose groups (**Fig 1c**.). The highest viral RNA load was observed in a ferret in the high dose group but there was a less consistent pattern of RNA detection which did not continue past day 7 pc. In the medium dose group, 4/6 ferrets were found to have detectable viral RNA in their rectal swabs between day 2 and 8 pc. No viral RNA was detected in any of the rectal swabs collected from the low dose group following challenge.

Viral RNA was detected at quantifiable levels in the bronchoalveolar lavage (BAL) of each ferret euthanised (scheduled) on day 3 pc from the high dose (1/1) and medium dose (1/1) groups (**Fig. 1d**). Viral RNA was detected but not quantified for ferrets across all three challenge groups at day 5 and 7 pc. There was no viral RNA detected in the BAL of any of the other ferrets after scheduled euthanasia. No viral RNA was detected in the blood of ferrets from any group taken, as scheduled, on days 2, 5, 8, 11 and 14 pc (data not shown).

Illumina sequencing of nasal wash RNA extracts showed little variation between the genome isolated at days 5 and 6 and the original sequence of the virus inoculated into the ferrets. Only one non-synonymous SNP was identified, in the day five pc for a ferret from the medium dose group; a T2152I mutation within the orf1ab polyprotein, no further timepoints were collected for this animal as it was euthanised at day 5 pc.

### Clinical signs

The normalised summed incidence of clinical scores for each group of ferrets is shown in **Fig 2a** and total summed scores are shown in **Table 2**. At day 9 pc all 3/3 ferrets in the high dose group showed reduced activity, a similar observation was made in the medium dose group but later on day 10 pc. Reduced activity was accompanied by ruffled fur, a sign that the ferrets were not grooming regularly. By day 13 pc ferrets in the medium dose group stopped showing signs of reduced activity and by day 15 pc the high dose groups stopped showing signs of reduced activity. Ferrets in the high dose group had the highest normalised cumulative clinical score (summed across all time points) (14.01), followed by the medium dose group (6.99) with sporadic instances recorded in the low dose group. No fever (> 39.9°C) was detected in any ferret, in any group (**Fig. 2b**); instead body temperature remained within the normal range. No weight loss was observed in any ferret in any group, below baseline; however, the SARS-CoV-2 infected ferrets failed to gain as much weight as the ferrets in the control (PBS) group, although this difference was not statistically significant (**Fig. 2c**).

**Table 2.**
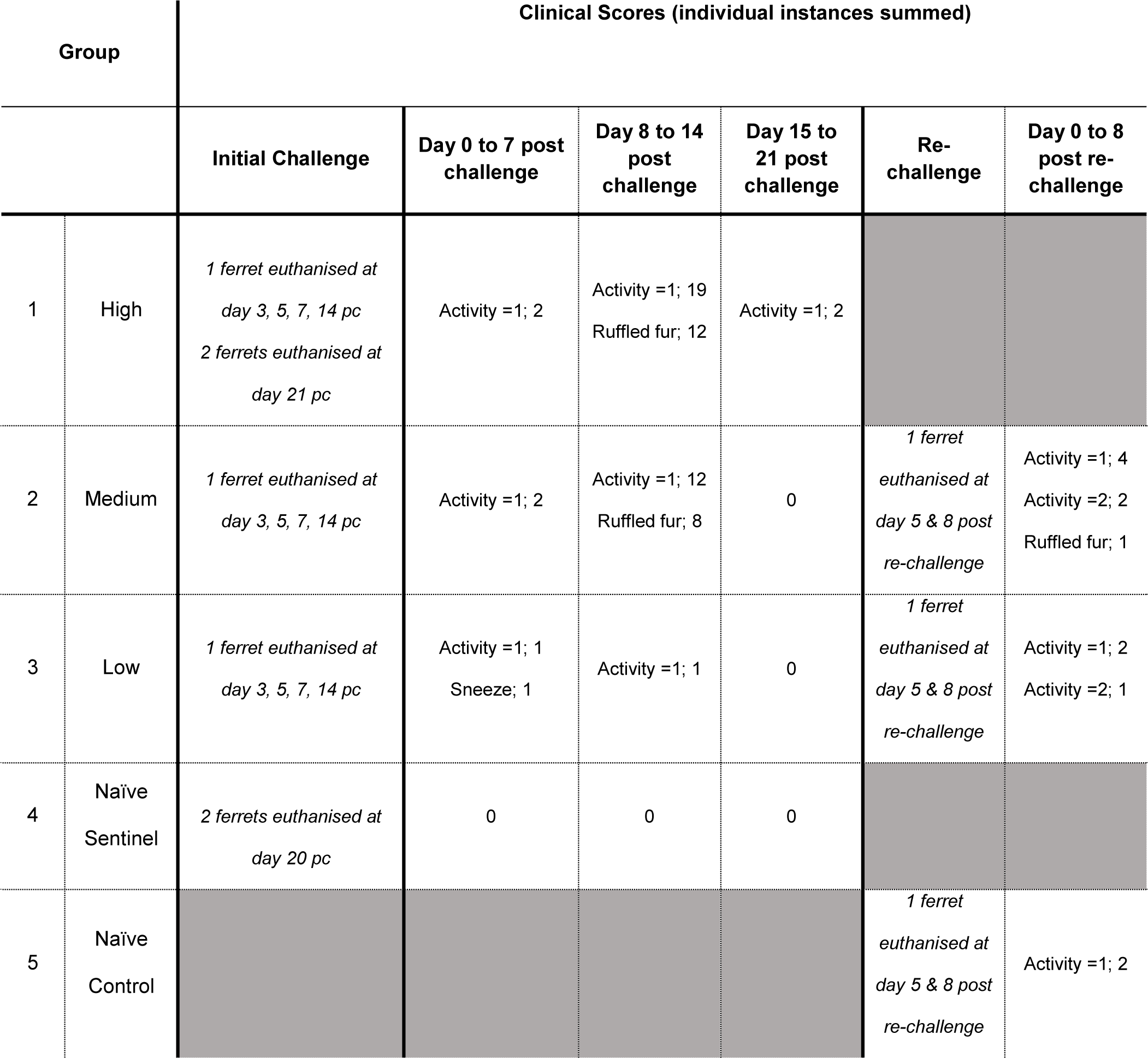
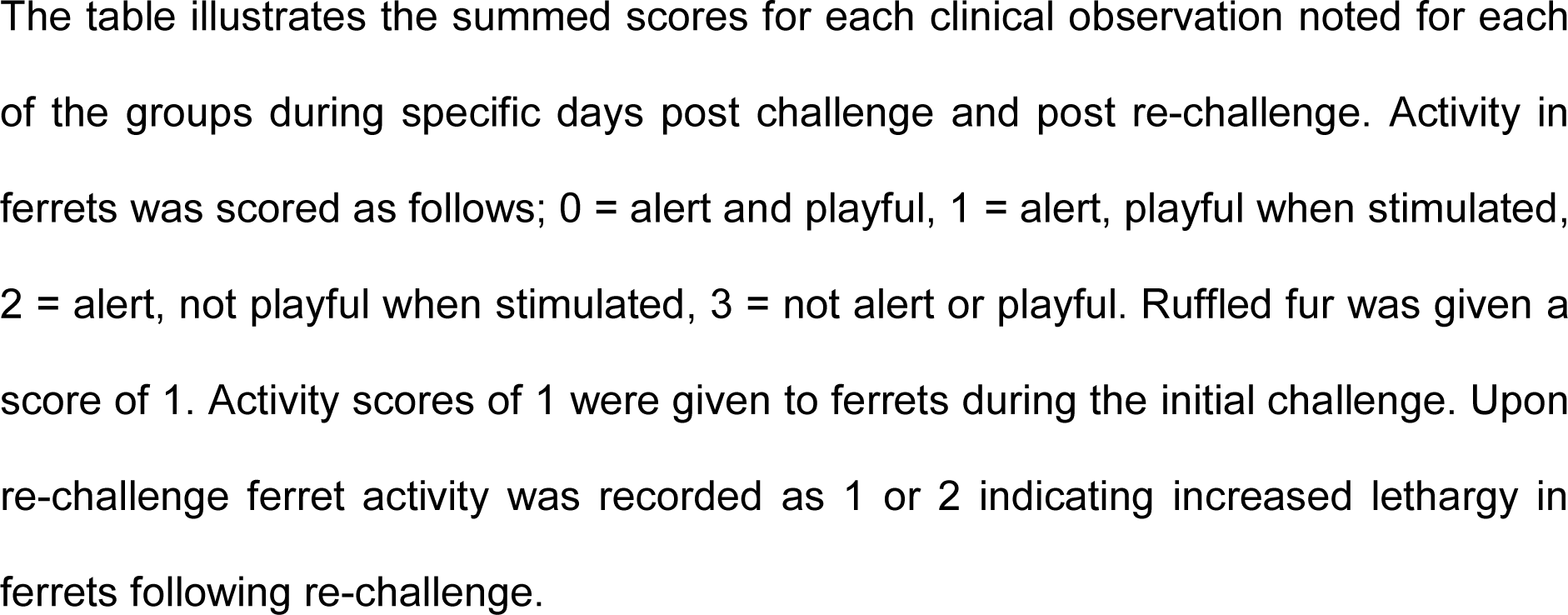
Clinical Scores.

| Group |  | Clinical Scores (individual instances summed) |  |  |  |  |  |
| --- | --- | --- | --- | --- | --- | --- | --- |
|  |  | Initial Challenge | Day 0 to 7 post challenge | Day 8 to 14 post challenge | Day 15 to 21 post challenge | Re-challenge | Day 0 to 8 post re-challenge |
| 1 | High | 1 ferret euthanised at day 3, 5, 7, 14 pc<br>2 ferrets euthanised at day 21 pc | Activity =1; 2 | Activity =1; 19<br>Ruffled fur; 12 | Activity =1; 2 |  |  |
| 2 | Medium | 1 ferret euthanised at day 3, 5, 7, 14 pc | Activity =1; 2 | Activity =1; 12<br>Ruffled fur; 8 | 0 | 1 ferret euthanised at day 5 & 8 post re-challenge | Activity =1; 4<br>Activity =2; 2<br>Ruffled fur; 1 |
| 3 | Low | 1 ferret euthanised at day 3, 5, 7, 14 pc | Activity =1; 1<br>Sneeze; 1 | Activity =1; 1 | 0 | 1 ferret euthanised at day 5 & 8 post re-challenge | Activity =1; 2<br>Activity =2; 1 |
| 4 | Naïve Sentinel | 2 ferrets euthanised at day 20 pc | 0 | 0 | 0 |  |  |
| 5 | Naïve Control |  |  |  |  | 1 ferret euthanised at day 5 & 8 post re-challenge | Activity =1; 2 |

The table illustrates the summed scores for each clinical observation noted for each of the groups during specific days post challenge and post re-challenge. Activity in ferrets was scored as follows; 0 = alert and playful, 1 = alert, playful when stimulated, 2 = alert, not playful when stimulated, 3 = not alert or playful. Ruffled fur was given a score of 1. Activity scores of 1 were given to ferrets during the initial challenge. Upon re-challenge ferret activity was recorded as 1 or 2 indicating increased lethargy in ferrets following re-challenge.

**Figure 2.**
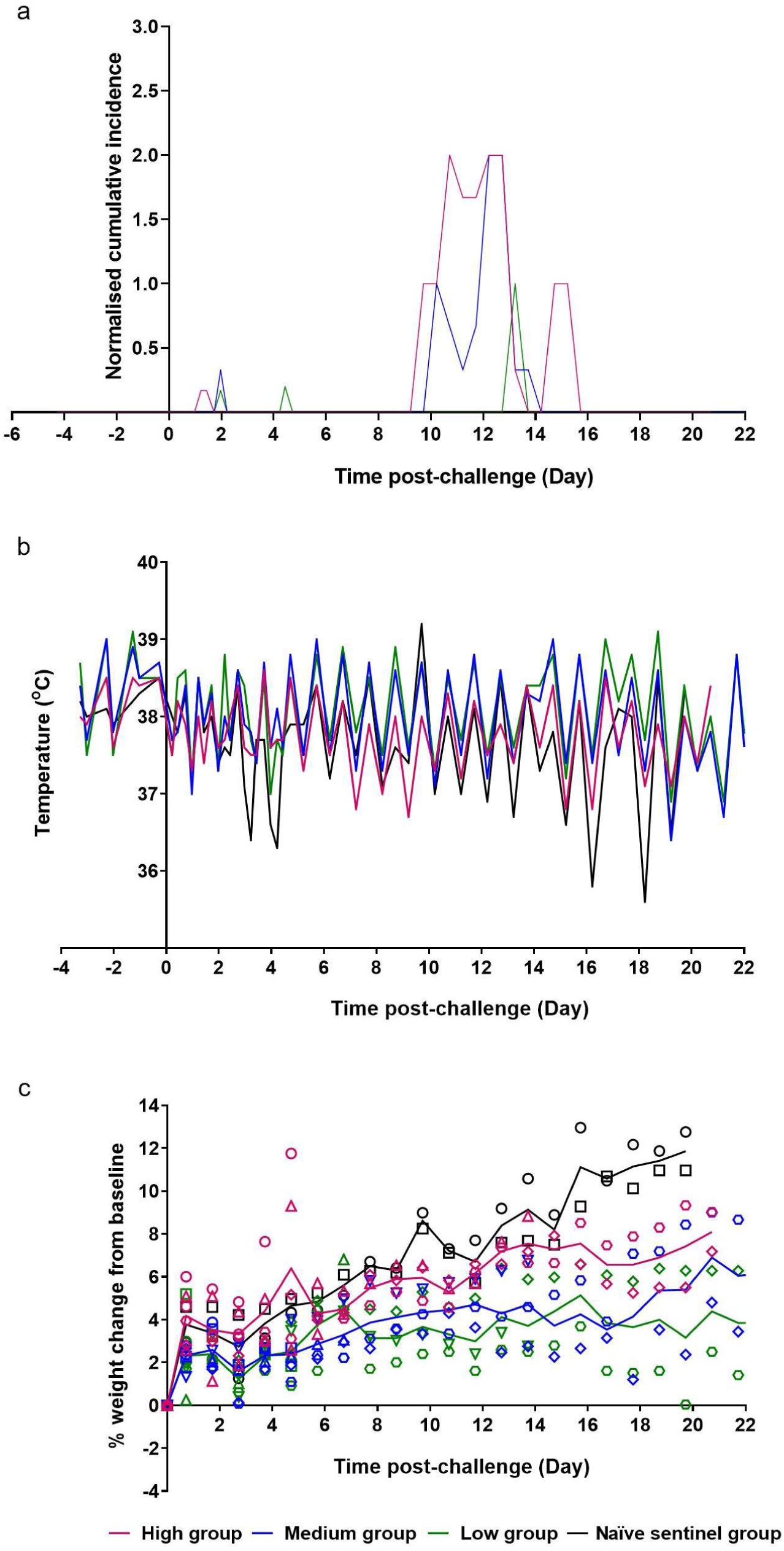
Clinical Observations. (**a**) Clinical observations were carried out four times daily (approximately 6 hours apart) for the first 5 days and then twice daily (approximately 8 hours apart) for the remaining time. Observations were summed for each group and normalised for the number of ferrets. (**b**) Temperatures were taken at the same time as clinical observations, using the identifier chip, to ensure any peak of fever was recorded. Mean temperatures are displayed on the graph. (**c**) Weight was recorded daily and percentage weight change from baseline was plotted. Points show values for individual animals, lines show group means.

### Histopathology

The nasal cavity from high dose ferrets showed a minimal to mild necrosis of epithelial cells (**Fig. 3a**) from days 3 to 7 pc. However, abundant epithelial cells from the nasal cavity were stained for viral RNA at day 3 pc (**Fig. 3b**). Occasional scattered cells expressing viral RNA were observed in high dose animals at days 5 and 7 pc and medium dose animals at days 3, 5 and 7 pc. Similarly, very few scattered epithelial cells were stained for viral RNA in the trachea and larynx from high and medium dose animals at day 3, 5 and 7 pc.

**Figure 3.**
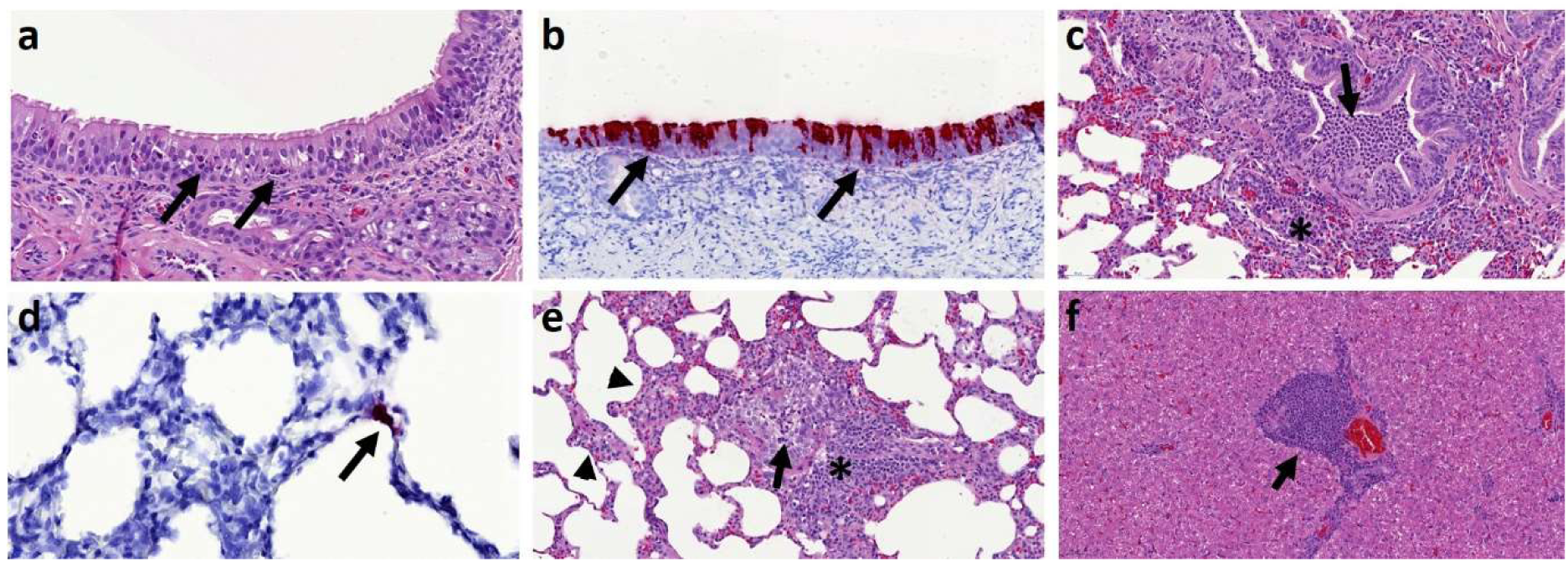
(**a**) Nasal cavity, day 3 pc, Group 1, H&E staining. Mild epithelial cell necrosis (arrows) and minimal inflammatory cell infiltration within the epithelium. (**b**) Nasal cavity, day 3 pc, Group 1, SARS-CoV-2 viral RNA detection (RNAScope staining). Presence of viral RNA in abundant ciliated epithelia cells from the nasal cavity mucosa. (**c**) Lung, day 5 pc, Group 1, H&E staining. Moderate bronchopneumonia with neutrophil and macrophage inflammatory infiltrate within the bronchiolar lumina (arrow). Mild peribronchiolar infiltration of mononuclear cells (*). (**d**) Lung, day 3 pc, Group 2, SARS-CoV-2 viral RNA detection (RNAScope staining). Presence of viral RNA in type II pneumocyte (arrow). (**e**) Lung, day 21 pc, Group 1, H&E staining. A Bronchiole with mild inflammatory infiltration in the lumina (arrow) and attenuation of the epithelial cells. Moderate peribronchiolar infiltration of mononuclear cells (*) and mild interalveolar septal inflammatory cell infiltration with thickening of the wall (arrowheads). (**f**) Liver, day 21 pc, Group 1, H&E staining. Moderate multifocal hepatitis with mononuclear cell infiltration in the portal areas (arrow).

No remarkable gross lesions were observed in the infected animals. Upon histological examination of the lungs of ferrets from the high and medium dose groups, a mild multifocal bronchopneumonia from day 3 to 14 pc was observed. Mild necrosis of the bronchiolar epithelial cells was observed together with inflammatory cell infiltration of neutrophils and mononuclear cells within the bronchiolar luminae, mostly affecting animals from the high dose group at day 3, 5 and 7 pc (**Fig. 3c**). This bronchopneumonia was characterised by the infiltration of inflammatory cells, mostly neutrophils, but also macrophages and lymphocytes, in approximately 10-15% of the lung section at day 3 pc decreasing to less than 5% at days 5 and 7 pc. The medium dose group showed mild bronchopneumonia in less than 5% of the lung sections at days 3 and 5 pc, while only occasional infiltration was observed in animals from the low dose group. Few cells stained positive for viral RNA using *in situ* hybridisation (RNAScope). Few type I and occasionally type II pneumocytes and alveolar macrophages were positive for viral RNA at days 3, 5 and 7 pc (**Fig. 3d**) in high and middle dose animals. Occasionally, mild proliferation of BALT was observed surrounding damaged bronchi and bronchioles at the early stages of the disease, with slightly more severity at day 14 and 21 pc (high dose) (**Fig. 3e**). Mild interstitial pneumonia with an increase in the thickness of the interalveolar septa was observed from day 3 pc towards the end of the experiment in high and medium dose groups (**Fig. 3e**). Ferrets from the high dose group showed mild proliferation of type II pneumocytes from day 7 pc onwards.

The liver showed multiple foci of inflammatory cell infiltration in the portal areas, composed of mainly macrophages, lymphocytes and occasional plasma cells (**Fig. 3f**). This multifocal infiltration was more severe in animals from the high and medium dose groups from day 3 pc, compared to the low dose group or control (PBS) animals, which only showed minimal presence of portal inflammation. No other remarkable changes were observed in any other tissue. However, occasional positive cells (absorbing epithelial enterocytes and goblet cells) were also observed in the small and large intestine from high and medium dose at days 3, 5 and 7 pc.

### Antibody Response to SARS-CoV-2 Infection

Neutralising antibody titres for ferrets infected in the high dose and medium dose groups generally increased longitudinally following challenge as illustrated in **Table 3**. The average fold increase of neutralising antibodies from day 8 pc to day 14 pc was about the same for both the high and medium dose groups. The low dose group had comparatively low neutralising antibodies throughout the time course. One naïve sentinel group ferret was shown to have strong neutralising antibodies to SARS-CoV-2 upon euthanasia at day 20 pc. This ferret showed no clinical signs of SARS-CoV-2 infection and samples takes at baseline, day 11 and day 20 post challenge were shown to be PCR negative for SARS-CoV-2. Further there was no evidence for any pathology in any of the tissues taken from both naïve sentinel ferrets euthanised on day 20 pc. Interestingly, the cellular immune response seen in lung mononucleocytes (MNCs) (data not shown) of the naïve sentinel ferret showed a high SARS-CoV-2 specific immune response to whole live SARS-CoV-2, paralleling the high neutralising antibodies seen in the plaque reduction neutralisation test for this ferret.

**Table 3.**
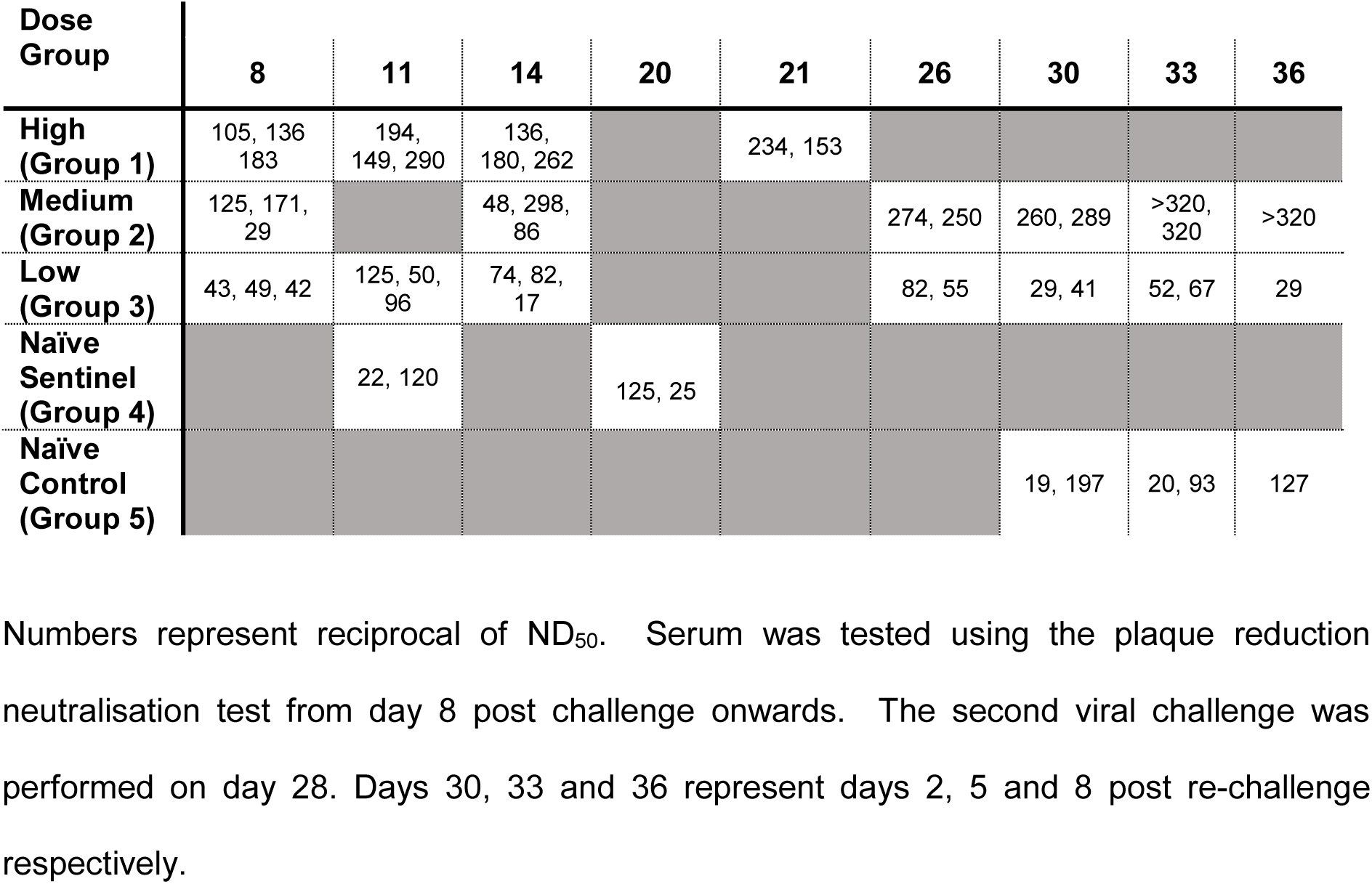
Neutralising Antibodies to SARS-CoV-2.

| Dose Group | 8 | 11 | 14 | 20 | 21 | 26 | 30 | 33 | 36 |
| --- | --- | --- | --- | --- | --- | --- | --- | --- | --- |
| High (Group 1) | 105, 136, 183 | 194, 149, 290 | 136, 180, 262 |  | 234, 153 |  |  |  |  |
| Medium (Group 2) | 125, 171, 29 |  | 48, 298, 86 |  |  | 274, 250 | 260, 289 | >320, 320 | >320 |
| Low (Group 3) | 43, 49, 42 | 125, 50, 96 | 74, 82, 17 |  |  | 82, 55 | 29, 41 | 52, 67 | 29 |
| Naïve Sentinel (Group 4) |  | 22, 120 |  | 125, 25 |  |  |  |  |  |
| Naïve Control (Group 5) |  |  |  |  |  |  | 19, 197 | 20, 93 | 127 |

Numbers represent reciprocal of ND<sub>50</sub>. Serum was tested using the plaque reduction neutralisation test from day 8 post challenge onwards. The second viral challenge was performed on day 28. Days 30, 33 and 36 represent days 2, 5 and 8 post re-challenge respectively.

### Re-challenge of ferrets with high dose SARS-Cov-2 results in absence of lung pathology

Four previously infected ferrets, two from the medium and low dose challenge groups, had neutralising titres of 1:274, 1:250, 1:82, and 1:55 at day 26 post challenge respectively, see **Table 3**. At day 28 pc, these ferrets and two naïve control animals were challenged intranasally with the high dose of SARS-Cov-2 (5×10^6^ pfu). Though URT infection was similar in all groups on day 2 post re-challenge (day 30 pc for Groups 1-3), viral RNA levels subsequently decreased in the previously challenged animals (n=4), with the medium dose group showing rapid decrease to below quantifiable levels by day 5 post re-challenge (day 33 pc). Viral RNA levels continued to stay above quantifiable levels in the naïve control group, although they began to fall at day 8 post re-challenge (**Fig. 4a**). Similar results were seen in the throat swab and rectal swabs (data not shown), with reduced viral shedding seen in the re-challenged animals. Animals in the re-challenged medium and low dose groups exhibited weight loss from baseline that was not seen at initial challenge for any of the animals in any of the challenge groups (**Fig. 4b**). Re-challenged animals also experienced increased clinical observations of lethargy and ruffled fur that was not observed at such an early stage in the initial challenge (**Table 2**). In contrast, the two previously naïve control animals did not experience weight loss below baseline after infection and they did not suffer the same level of clinical observation as the re-challenged animals (**Fig 4b**).

**Figure 4.**
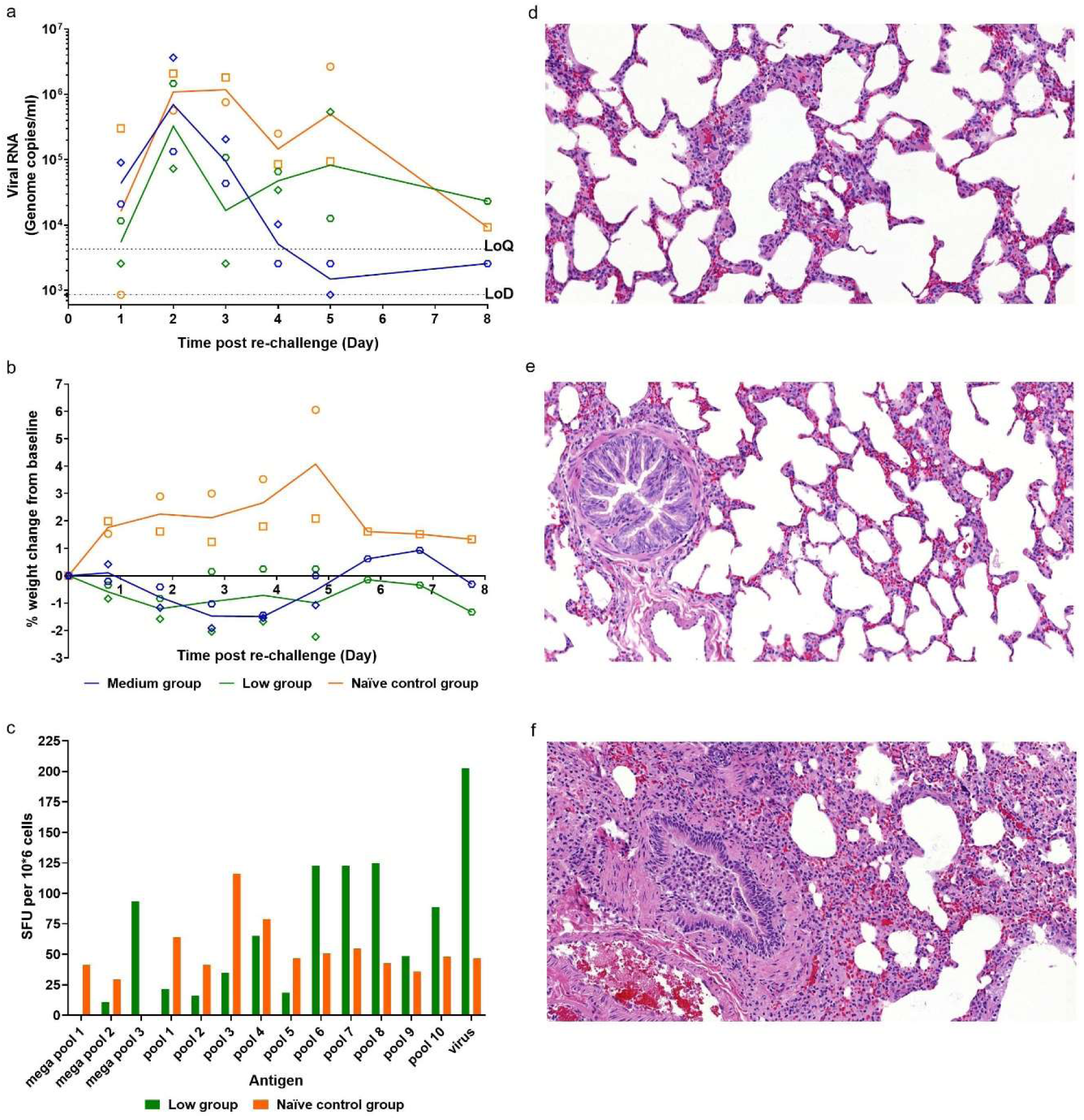
Re-challenge of ferrets with SARS-CoV-2. (**a**) Nasal washes were collected at days 1-5 post re-challenge (days 29 – 33 post-original challenge). Viral RNA was quantified by RT-qPCR. (**b**) Percentage weight change from baseline. Baseline was calculated as average of the two most recent weights taken preceding re-challenge. (**c**) Cellular immune responses of ferrets infected with SARS-CoV-2. Lung MNCs were collected from animals (n= 1) at day 36 pc (day 8 following re-challenge). SARS-CoV-2 specific IFN-γ responses were seen in both infected ferrets. The values measured for each ferret are plotted as spot forming units (SFU) per million cells. (**d**) Medium dose re-challenged ferret at day 5 post re-challenge. No remarkable changes in alveoli or terminal bronchiole. (**e**) Low dose re-challenged ferret at day 5 post re-challenge. No remarkable changes in alveoli or bronchiole. (**f**) Control group ferret challenged for the first time (day 28 pc). Inflammatory infiltration within bronchiolar lumen and mild infiltration of alveolar septa; lesions comparable with those observed in the original high dose group.

The cellular immune response in the lungs of a low dose (Group 2) re-challenge ferret and a naïve control (Group 5) ferret at day 36 (8 days post re-challenge respectively) were compared. **Fig. 4c** shows SARS-CoV-2 specific cellular immune responses, as determined by IFN-γ ELISpot. The number of secreting cells detected after re-stimulation of lung MNCs with peptide pools spanning the spike protein varied between ferrets from each group. The strongest response is detected in the re-challenge ferret after ex-vivo re-stimulation with whole live virus. Upon histological examination the upper and lower respiratory tracts from animals in both re-challenged groups showed no remarkable lesions (**Fig. 4d and 4e**), and no presence of the significant bronchopneumonia that was observed in ferrets challenged with 5×10^6^ pfu for the first time, i.e. the original high dose ferret or the naïve control infected group included for the ‘re-challenge’ (**Fig. 4f**). This parallels the absence of pathology observed in the two naïve sentinel ferrets euthanised at day 20 pc.

## Discussion

This study demonstrated that ferrets are susceptible to experimental intranasal infection with a low passage isolate of SARS-CoV-2 strain Victoria 1. A high dose (5×10^6^ pfu/ml in a 1ml volume) intranasal challenge in ferrets produced mild clinical signs, consistent lung pathology and a viral shedding pattern that aligns with the mild to moderate disease seen in the human population.

Previously published SARS-CoV-1 challenge studies conducted in the ferret show that a lower dose of that virus (10^3^ TCID_50_) is sufficient to cause a mild disease in the ferret ^20,21^. Here it was shown that a high (5×10^6^ pfu) and medium (5×10^4^ pfu) dose intranasal challenge resulted in an infection characterised by prolonged viral RNA shedding in all ferrets (days 1 -11 pc), accompanied by observable clinical signs from day 8 post challenge for both high and medium dose groups. Onset of clinical symptoms were delayed by approximately 24 hours in the medium dose animals. Both doses also induced classical pathology of bronchial pneumonia involving 10% and 3% of recipient lungs respectively. A low dose intranasal challenge of the same SARS-CoV-2 virus (5×10^2^ pfu) appeared to result in infection of only one ferret which shed viral RNA in the upper respiratory tract and failed to show any remarkable lesions in the respiratory tract.

In the high and medium dose groups, virus was readily detected using *in-situ* hybridisation in the upper respiratory tract of ferrets, with a peak at 3 days post challenge. These findings aligned with the detected shedding of viral RNA from nasal washes which also peaked at day 3 to 4 post challenge. This upper respiratory infection mirrors the clinical disease recently reported in mild cases of humans infected with SARS-CoV-2 infection^28^.

Recent reports indicate that COVID-19 patients appear to shed viral RNA intermittently after recovery from disease with some individuals being tested and found to be positive again after being released from isolation^29^. This report is in alignment with the observations in the two ferrets challenged with the high dose of SARS-CoV-2 (euthanised at day 21) which appeared to continue to shed detectable viral RNA from the upper respiratory tract up to day 18 post challenge even though these animals had developed neutralising antibodies.

The main histopathological finding in approximately 10% of the lung tissue sections in the high dose group consisted mainly of a multifocal bronchiolitis, with inflammatory infiltrates within the airways and some alveolar species. This finding is similar, but less severe, to the findings in the published reports about SARS-CoV-1 ferret challenge models^22,30,31^. In this study, mild alveolar damage was observed in the acute phase. At later time points, mild proliferation of type II pneumocytes, with interstitial infiltrates and peribronchiolar cuffing, was recoded, consistent with evolution form the acute phase.

Mild to moderate multifocal hepatic inflammatory cell infiltration has been widely reported in viral infections in animals, and has been previously described in SARS-CoV-1 infected ferrets^32^. However, the periportal infiltrates may not be associated with injury to the surrounding tissue and they are reported as a common background finding in laboratory ferret species. The presence of infected enterocytes has been reported for SARS-CoV-1 and SARS-CoV-2 in humans^33^ and different ferret models^34,35^.

The upper respiratory virus replication, reported here, in the high and medium dose groups of animals, support the observations of Shi *et al*. ^34^ who challenged ferrets with 10^5^ pfu of SARS-CoV-2 and found peak levels of viral RNA (10^8^ copies/ml) in nasal washes on day 6. They also reported mild lung pathology associated with SARS-CoV-2 infection similar to our medium dose animals, but this was not as extensive as that seen in our high dose challenge group ferrets.

A ferret model of SARS-CoV-2 infection has also been developed by Kim *et al*.^35^ who used a challenge of 3×10^5^ pfu which resulted in peak URT viral RNA shedding on day 4-6 which coincided with a significant temperature spike. In the present study, animals challenged with either 5×10^4^ or 5×10^6^ pfu displayed a consistent fatigue after peak viral shedding, however a significant increase in body temperature was not observed. This fatigue after peak of viral shedding was not observed in the low dose group, ruling out the possibility of it being induced by an aspect of ferret handling, such as sedation or sampling.

Both Shi *et al*.^34^ and Kim *et al*.^35^ report live virus isolation from RNA positive nasal wash samples. In this study, only low levels of live virus in some nasal washes and throat swabs were detected even though high levels of viral RNA were detected. A possible reason for this observation could be poor stability of this virus isolate, which is currently untested, resulting in a knockdown of live virus between taking the samples from a ferret and assaying the material. There may also have been an inhibitory effect from the sample matrix in the cell cultures used for the live virus plaque assay. Alternatively, this result may have accurately reflected low levels of viable virus presence which others have reported even though viral RNA can be detected. For human swabs and sputum samples, it has been noted that infectious virus was never recovered from samples with a viral RNA load of less than 10^6^ copies/ml^28^.

Neutralising antibody levels developed in ferrets in all challenge groups within 14 days, even though some animals in the low dose group were found to have no detectable viral RNA shedding. The finding that low and medium dosed animals showed reduced viral RNA shedding in the URT and an absence of lung pathology following re-challenge is encouraging; it suggests that there may be potential benefits for the healthy population as a result of naturally acquired immunity and is in line with the observation reported by Bao *et al*. ^36^ in which previously infected rhesus macaques were protected against re-challenge with SARS-CoV-2.

SARS-CoV-2 spike protein-specific immune responses seen in a low dose re-challenged ferret were compared to that of a primary challenge ferret. This comparison showed that the response to the virus appears to be higher on re-challenge. However, ferrets challenged with our high dose of SARS-CoV-2 displayed increased clinical observations and lost weight from baseline following re-challenge, hinting at enhanced disease but a larger study would be required to effectively assess this observation. Alternatively, these clinical signs may be perfectly normal host response to infection in a pre-immune individual whilst the immune system is successfully clearing a large challenge dose.

This study demonstrated that ferrets challenged with 5×10^6^ pfu or 5×10^4^ pfu displayed only mild clinical signs of SARS-CoV-2 infection. These signs appeared to be less severe than those reported after ferrets were infected with SARS-CoV-1^21,30^.

This ferret model of intranasal SARS-CoV-2 infection presents three key measurable endpoints: a) consistent URT viral RNA shedding; b) significant lung pathology; and c) post viral fatigue. Reductions in URT RNA shedding during the first 14 days post intranasal challenge could be an attractive indicator of the efficacy of candidate therapeutics and vaccines. It may be wise, however, to euthanise prior to 14 days post challenge to more accurately assess the impact on lung pathology especially when looking for signs of vaccine-enhanced disease^20,37,38^. Alternatively, if CT scanning facilities were available, this may be achieved without the need to euthanise a cohort if this technology was found suitable to also make this assessment. We believe the high dose intranasal challenge will provide the most distinct disease endpoints. However, with its reduced level of lung pathology, the medium dose challenge may provide a higher level of sensitivity to some interventions, as was observed when assessing therapeutics to influenza in the ferret model^39^.

## Materials & Methods

### Viruses and Cells

SARS-CoV-2 Victoria/01/2020^26^ was generously provided by The Doherty Institute, Melbourne, Australia at P1 and passaged twice in Vero/hSLAM cells [ECACC 04091501] Whole genome sequencing was performed, on the challenge isolate, using both Nanopore and Illumina as described previously^40^. Virus titre was determined by plaque assay on Vero/E6 cells [ECACC 85020206]. Cell lines were obtained from the European Collection of Authenticated Cell Cultures (ECACC) PHE, Porton Down, UK. Cell cultures were maintained at 37oC in MEM (Life Technologies, California, USA) supplemented with 10% foetal bovine serum (Sigma, Dorset, UK) and 25 mM HEPES (Life Technologies).

### Animals

Twenty-two healthy, female ferrets (*Mustela putorius furo*) ages 7 months were obtained from a UK Home Office accredited supplier (Highgate Farm, UK). The mean weight at the time of challenge was 1032g/ferret (range 870-1239g). Animals were housed in pairs at Advisory Committee on Dangerous Pathogens (ACDP) containment level 3. Cages met with the UK Home Office *Code of Practice for the Housing and Care of Animals Bred, Supplied or Used for Scientific Procedures* (December 2014). Access to food and water was *ad libitum* and environmental enrichment was provided. All experimental work was conducted under the authority of a UK Home Office approved project licence that had been subject to local ethical review at PHE Porton Down by the Animal Welfare and Ethical Review Body (AWERB) as required by the *Home Office Animals (Scientific Procedures) Act 1986*.

### Experimental Design

Before the start of the experiment animals were randomly assigned to challenge groups, to minimise bias. The weight distribution of the animals was tested to ensure there was no statistically significant difference between groups (one-way ANOVA, p > 0.05). An identifier chip (Bio-Thermo Identichip, Animalcare Ltd, UK) was inserted subcutaneously into the dorsal cervical region of each animal. Prior to challenge animals were sedated by intramuscular injection of ketamine/xylazine (17.9 mg/kg and 3.6 mg/kg bodyweight). Challenge virus was delivered by intranasal instillation (1.0 ml total, 0.5 ml per nostril) diluted in phosphate buffered saline (PBS).

Three different doses of virus were delivered to three groups (n=6) of ferrets: high [5×10^6^ pfu/ml], medium [5×10^4^ pfu/ml] and a low [5×10^2^ pfu/ml] dose. For the high, medium and low dose groups, individual ferrets were scheduled for euthanasia on day 3 (n=1), day 5 (n=1), day 7 (n=1) and day 14 (n=1). For the high dose group, the remaining 2 ferrets were euthanised on day 21 (n=2). The mock-infected animals (n=2) received an intranasal instillation of sterile PBS and were euthanised on day 20.

On day 28 pc the remaining ferrets in the low (n=2) and medium (n=2) groups were re-challenged with 5×10^6^ pfu by the intranasal route. Additional naïve control ferrets (n=2) were also challenged on day 28, to provide a re-challenge control. All 6 animals were monitored for clinical signs and one ferret from each group was euthanised on day 33 and the remaining animals were euthanised on day 36.

Nasal washes, throat and rectal swabs were taken at days -1, 1-8, 10, 11, 13, 14, 16, 18 and 20 pc. They were also taken at days 1-5 and 8 post re challenge (days 29-33 and 36 pc). Whole blood and serum were collected at 2, 5, 8, 11, 14 days pc for all ferrets. Whole blood and serum were collected at days 2, 5 and 8 (days 30, 33 and 36 pc) post re-challenge for all remaining ferrets. The negative control ferrets (n=2) had nasal washes, throat swabs, whole blood and serum taken at -1 and 11 days pc. At necropsy nasal washes, throat and rectal swabs, whole blood and serum were taken alongside tissue samples for histopathology. Nasal washes were obtained by flushing the nasal cavity with 2 ml PBS. For throat swabs, a flocked swab (MWE Medical Wire, Corsham, UK) was gently stroked across the back of the pharynx in the tonsillar area. Throat and rectal swabs were processed, and aliquots stored in viral transport media (VTM) and AVL at -80C until assay.

### Clinical and euthanasia observations

Animals were monitored for clinical signs of disease four times daily (approximately 6 hours apart) for the first 5 days pc and then twice daily (approximately 8 hours apart) for the remaining time. Clinical signs of disease were assigned a score based upon the following criteria. Activity was scored as follows; 0 = alert and playful, 1 = alert, playful when stimulated, 2 = alert, not playful when stimulated, 3 = not alert or playful. Ruffled fur was given a score of 1. No other clinical signs were noted. In order to meet the requirement of the project license, immobility, neurological signs or a sudden drop in temperature were automatic euthanasia criteria. Animals were also deemed to have reached a humane endpoint if their body weight was at or below 30% baseline. If any ferret reached any of these three euthanasia criteria, they were to be immediately euthanised using a UK Home Office approved Schedule 1 procedure. However, no animals reached these end-points during this study.

Temperature was taken using a microchip reader and implanted temperature/ID chip. Temperature was recorded at each clinical scoring point using the chip to ensure any peak of fever was recorded. Animals were weighed at the same time of each day from the day before infection until euthanasia.

### Necropsy Procedures

Ferrets were anaesthetised with ketamine/xylazine (17.9 mg/kg and 3.6 mg/kg bodyweight) and exsanguination was effected via cardiac puncture, followed by injection of an anaesthetic overdose (sodium pentabarbitone Dolelethal, Vetquinol UK Ltd, 140 mg/kg). A necropsy was performed immediately after confirmation of death. The bronchoalveolar lavage (BAL) was collected at necropsy from the right lung. The left lung was dissected prior to BAL collection and used for subsequent histopathology and virology procedures.

### RNA Extraction

RNA was extracted using a Viral RNA QIAamp kit (Qiagen) following manufacturer’s instruction. RNA was isolated from nasal wash, throat and rectal swabs, EDTA treated whole blood and BAL.

### Quantification of Viral Loads by RT-qPCR

Reverse transcription-quantitative polymerase chain reaction (RT-qPCR) targeting a region of the SARS-CoV-2 nucleocapsid (N) gene was used to determine viral loads and was performed using TaqPath™ 1-Step RT-qPCR Master Mix, CG (Applied Biosystems™), 2019-nCoV CDC RUO Kit (Integrated DNA Technologies) and 7500 Fast Real-Time PCR System (Applied Biosystems™) was used. Sequences of the N1 primers and probe were: 2019-nCoV_N1-forward, 5’ GACCCCAAAATCAGCGAAAT 3’; 2019-nCoV_N1-reverse, 5’ TCTGGTTACTGCCAGTTGAATCTG 3’; 2019-nCoV_N1-probe, 5’ FAM-ACCCCGCATTACGTTTGGTGGACC-BHQ1 3’. The cycling conditions were: 25°C for 2 minutes, 50°C for 15 minutes, 95°C for 2 minutes, followed by 45 cycles of 95°C for 3 seconds, 55°C for 30 seconds. The quantification standard was a 100bp Ultramer RNA oligo (Integrated DNA Technologies) equivalent to 28274-28373bp of SARS-CoV-2 NC_045512.2, with quantification between 1 × 10^1^ and 1 × 10^7^ copies/µl. Positive samples detected below the limit of quantification were assigned the value of 6 copies/µl, whilst undetected samples were assigned the value of ≤ 2 copies/µl, equivalent to the assays limit of detection.

### SARS-CoV-2 virus plaque assay

Samples were diluted in serum-free MEM containing antibiotic/antimycotic (Life Technologies) and incubated in 24-well plates (Nunc, ThermoFisher Scientific, Loughborough, UK) with Vero E6 cell monolayers. Virus was allowed to adsorb at 37 °C for 1 hour, then overlaid with MEM containing 1.5% carboxymethylcellulose (Sigma), 4% (v/v) foetal bovine serum (Sigma) and 25 mM HEPES buffer (Life Technologies). After incubation at 37 °C for 5 days, they were fixed overnight with 20% (w/v) formalin/PBS, washed with tap water and stained with methyl crystal violet solution (0.2% v/v) (Sigma).

### Plaque Reduction Neutralisation Test

Neutralising virus titres were measured in heat-inactivated (56°C for 30 min) serum samples. SARS-CoV-2 was diluted to a concentration of 933 pfu/ml (70 pfu/75 µl) and mixed 50:50 in 1% FCS/MEM with doubling serum dilutions from 1:10 to 1:320 in a 96-well V-bottomed plate. The plate was incubated at 37°C in a humidified box for 1 hour to allow the antibody in the serum samples to neutralise the virus. The neutralised virus was transferred into the wells of a washed plaque assay 24-well plate (see plaque assay method), allowed to adsorb at 37°C for a further hour, and overlaid with plaque assay overlay media. After 5 days incubation at 37°C in a humified box, the plates were fixed, stained and plaques counted. Median neutralising titres (ND_50_) were determined using the Spearman-Karber ^41^ formula relative to virus only control wells.

### Histopathological Analysis

Samples from the left cranial and left caudal lung lobe together with spleen, kidney, liver, tracheobronchial and axillary lymph nodes, jejunum, colon, trachea, larynx and nasal cavity, were fixed by immersion in 10% neutral-buffered formalin and processed routinely into paraffin wax. Nasal cavity samples were decalcified using an EDTA-based solution prior to embedding. 4 µm sections were cut and stained with haematoxylin and eosin (H&E) and examined microscopically. In addition, samples were stained using the RNAscope technique to identify the SARS-CoV-2 virus RNA. Briefly, tissues were pre-treated with hydrogen peroxide for 10 minutes (room temperature), target retrieval for 15 mins (98-101°C) and protease plus for 30 mins (40°C) (Advanced Cell Diagnostics). A V-nCoV2019-S probe (Cat No. 848561, Advanced Cell Diagnostics) was incubated on the tissues for 2 hours at 40°C. Amplification of the signal was carried out following the RNAscope protocol using the RNAscope 2.5 HD Detection kit – Red (Advanced Cell Diagnostics).

### Isolation of Lung Mononuclear Cells

Whole lungs were removed from each ferret. The lungs were dissected into small pieces and placed into a 12.5ml solution of collagenase (715 collagenase units/ml) (Sigma-Aldrich) and DNase (350 DNase units/ml) (Sigma-Aldrich). Lungs were placed into gentleMACS C-tubes and agitated whilst incubating at 100rpm, 37oC for 1 hour on an OctoMACS (Miltenyi Biotec, Surrey, UK). Partially digested lung tissue was then dissociated using an OctoMACS. The tissue solution was passed through two cell sieves (100µm then 70µm) and then layered with Ficoll^®^-Paque Premium (GE Healthcare, Hatfield, United Kingdom). Density gradient centrifugation was carried out at 400g for 30 minutes. Buffy coats containing lymphocytes were collected and washed with medium by pelleting cells via centrifugation at 400 g for 10 minutes. The cells were counted using a vial-1 cassette and a Nucleocounter-200 before cryopreservation in 95% FCS/ 5% v/v DMSO. Cryopreserved cells were then frozen at -80°C in controlled rate freezer containers overnight, before transfer to liquid nitrogen (vapour phase).

### Interferon-gamma (IFN-γ) ELISpot Assay

An IFN-γ ELISpot assay was performed to determine the production capacity of SARS-CoV-2-specific T cells in the lung using a ferret IFN-γ kit (Mab-tech, Nacka. Sweden). Lung MNCs were defrosted into pre-warmed medium (R10) consisting of RPMI 1640 medium (Sigma-Aldrich) supplemented with 2mM L-glutamine (Sigma-Aldrich), 0.05mM 2-mercaptoethanol (Invitrogen, Paisley, United Kingdom), 25mM HEPES buffer (Sigma-Aldrich, Dorset, United Kingdom), 100 U/ml Penicillin/100 µg/ml Streptomycin solution (Sigma-Aldrich), 10% heat inactivated foetal bovine serum (Sigma-Aldrich), and benzonase (Novogen, Merck, Darmstadt, Germany). Cells were rested for 2 hours prior to use. Lung MNCs were assessed for responses to whole SARS-CoV-2 virus and a COVID-19 Spike Protein (GenBank: QHQ82464.1) peptide panel. The peptide panel consisted of 15mer peptides overlapping by 11mers. Individual peptides were reconstituted in 10% v/v DMSO. The 10 peptide pools, each containing 32 peptides, were created by combining equimolar amounts of each peptide. Three mega pools spanning the whole spike protein (approx. 100 peptides in each mega pool) were also created. Each peptide pool and mega pool was diluted for use in the ELISpot assay in supplemented RPMI to achieve a final concentration of at 2.5µg per peptide. SARS-CoV-2 whole virus was also used at an MOI of 0.09 to re-stimulate the lung MNCs. The virus was cell culture grown and was a direct match to the isolate used for ferret challenge. R10 media was used as a negative control and for preparation and dilution of cells, peptide, virus and stimulants. Cell stimulation cocktail (PMA/Ionomycin 500x concentrate, Sigma-Aldrich, Dorset, United Kingdom), was used as a positive control to prove cells were capable of a stimulation response. Pre-coated ferret anti-IFN-γ ELISpot plates (mAb MTF14, Mab-tech, Nacka. Sweden) were used and 500,000 lung MNCs were plated per well in 50µl of R10, with or without antigen, in duplicate and incubated overnight (37°C, 5% CO_2_). Following cell stimulation, plates were washed 5x with 1x PBS (Gibco) and incubated at RT for 2 hours with biotinylated anti IFN-γ IgG. The plates were then washed 5x with 1x PBS and incubated with streptavidin-ALP for 1 hour, RT. The plates were washed again 5x with 1x PBS and spots were developed with 5-bromo-4-chloro-3-indoly phosphate (BCIP)-Nitro Blue tetrazolium (NBT) substrate. Plates were allowed to dry overnight and decontaminated by formaldehyde fumigation before removal from the CL3 facility. Plates were read, counted and quality control checked using the CTL ELISpot plate reader and ImmunoSpot 5.0 analyser software. Results from duplicate test wells were averaged. Data were corrected for background by subtracting the mean number of spots from the R10 media control wells from the mean counts of spots in the test wells.

## Acknowledgements

The authors gratefully acknowledge the support from the Biological Investigations Group at the National Infection Service, PHE, Porton Down, United Kingdom. The views expressed in this paper are those of the authors and not necessarily those of the funding body. This work was funded by the US Food and Drug Administration [Contract number: HHSF223201710194C].

## Contributions

K.A.R, Y.H, S.G.F, C.J.W, J.A.H and M.W.C. conceived the study.

J.D. and M.C. provided virus strain.

K.R.B. grew viral stock, optimised virology techniques and supervised virology experiments.

S.A.F, D.J.H, I.T. and N.R.W performed all animal procedures at containment level 3.

K.A.R, P.B, B.E.C, K.G, C.M.K.H, D.N, K.S and S.T processed all animal samples at containment level 3.

L.H, C.L.K, E.R and F.J.S contributed to histology experiments and performed critical assessment of pathology.

L.A, E.B, K.R.B, N.S.C, K.J.G, H.E.H, S.L and E.J.P contributed wet virology experiments and analysis of data.

K.A.R, L.A, E.B, J.G. R.H, S.L J.P, K.S and N.I.W contributed to inactivation, extraction and PCR of samples

G.S performed quality control and analytical assistance on PCR data

D.P.C, S.T.P and K.L.O performed NGS and analysis data.

K.A.R performed analysis on data generated

T.T, R.W and M.J.D provided technical assistance

K.A.R, J.A.T, F.J.S and M.W.C wrote the manuscript

A.C.M, C.J.W, K.S, B.E.C and S.G.F. provided critical review

